# Equine strongyle communities are constrained by horse sex and species dipersal-fecundity trade-off

**DOI:** 10.1101/225417

**Authors:** Sallé Guillaume, Kornaś Sławomir, Basiaga Marta

**Affiliations:** UMR1282 INRA/Université de Tours Infectiologie et Santé Publique, F-37380; Department of Environmental Zoology, Institute of Animal Sciences, University of Agriculture in Krakow, al. Mickiewicza 24/28, 30-059 Krakow, Poland

**Keywords:** horse, cyathostomin, strongyle, ecology, sex, nematode

## Abstract

Equine strongyles are a major health issue. Large strongyles can cause death of horses while cyathostomins (small strongyles) have shown increased resistance to anthelmintics worldwide. Description of strongyle communities have accumulated but little is known about the diversity of these communities and underpinning environmental factors.

This study analysed the diversity of strongyle communities in 48 horses from Poland. Strongyle species fell into two groups, contrasted by their prevalence and relative abundance. Seven horses were necessary to sample at least 90% of strongyle community diversity, providing a minimal cut-off to implement sampling trial in the field. Strongyle communities entertained a network of mostly positive interactions and species co-occurrence was found more often than expected by chance. In addition, species fecundity and prevalence were negatively correlated *r*=-0.78), suggesting functional trade-offs between species dispersal abilities and fecundity. This functional trade-off may underpin species coexistence. Horse sex was also a significant constraint shaping strongyle communities. Indeed, mares generally displayed more similar strongyle communities than stallions (p=0.004) and *Cylicostephanus calicatus* was more abundant in stallions suggesting sex-specific interactions (p=0.02). While niche partitioning is likely to explain some of the positive interactions between equine strongyle species, coexistence may also result from a functional trade-off between dispersal ability and fecundity. There is significant evidence that horse sex drives strongylid community structure, which may require differential control strategies between mares and stallions.

## 1. Introduction

Grazing horses are infected by a wide variety of intestinal strongyle parasites belonging to the Strongylinae and Cyathostominae subfamilies (Lichtenfels et al., 2008). Among the Strongylinae, *Strongylus vulgaris* has been identified as a major cause of colic in horses (Duncan and Campbell, 1973). This results from the accumulation of larval stages in the cranial mesenteric artery that leads to an arteritis responsible for fatal intestinal infarction (Duncan and Campbell, 1973). But modern anthelmintic treatments have dramatically decreased its prevalence (Nielsen et al., 2012). Cyathostominae (cyathostomin or small strongyles) form a wider tribe than equine Strongylinae, encompassing 51 species, 40 of which infecting horses (Lichtenfels et al., 2008). Unlike Strongylinae, these nematodes can reach prevalence up to 100% in horses (Lyons et al., 1999). Cyathostomins have been associated with milder clinical signs like poor hair coat and weight loss (Murphy and Love, 1997). Although, the massive emergence of larval stages from the colonic mucosa where they reside can lead to cyathostominosis cases, characterised by diarrhoea, protein-losing enteropathy (Love et al., 1999). Failure to control larval cyathostominosis can also lead to the death of affected horses in at least a third of cases, as suggested by a report on 15 clinical cases in the United-Kingdom (Giles et al., 1985). More importantly, cyathostomes have been involved in multiple reports of anthelmintic failures throughout the world thus representing a major issue in equine medicine (Traversa et al., 2012; Canever et al., 2013; Geurden et al., 2014; Relf et al., 2014).

Although numerous cyathostomins species have been described, necropsy reports consistently described cyathostomins communities as a core assemblage of ten to 13 species (Ogbourne, 1976; Mfitilodze and Hutchinson, 1990; Lyons et al., 1999; Collobert-Laugier et al., 2002; Kuzmina et al., 2005; Traversa et al., 2010). Notably, these reports also underscore the cyathostomin ability to colonize various ecological niches worldwide, being highly prevalent under tropical (Mfitilodze and Hutchinson, 1990; Silva et al., 1999), continental (Kuzmina et al., 2005; Kuzmina and Kharchenko, 2008) or oceanic and Mediterranean climatic conditions (Traversa et al., 2010).

Despite the wealth of reports describing equine strongyle communities, scant knowledge has been gathered about the drivers underpinning strongyle biodiversity or community assemblage. Nonetheless, some efforts have been made to characterise the consequences of deworming treatments on strongyle diversity as an attempt to better understand widespread anthelmintic failures. First, a reduced diversity in strongyle communities was observed in Ukrainian farms applying frequent deworming (Kuzmina and Kharchenko, 2008). Second, investigations of strongyle community structure in horse populations showing reduced egg reappearance period after anthelmintic treatment, suggested that a limited number of species (*Cylicocyclus nassatus* or *Cylicostephanus longibursatus*) was driving the drug resistance phenotype (van Doorn et al., 2014; Kooyman et al., 2016). However, these observations might be mixed up with the reported variation in strongyle species fecundity (Kuzmina et al., 2012), dispersal ability, or relative abundance. Indeed, these latter properties may affect the post-treatment community structure, *e.g*. the most abundant species contributing most to the post-treatment community, without any genuine association with anthelmintic susceptibility. Some insights into the structure of equine strongyle communities have been provided by a report focusing on pair-wise correlations between strongyle species abundance (Stancampiano et al., 2010). This analysis revealed negative interactions between major species, suggesting competition might be involved (Stancampiano et al., 2010). In addition, an attempt to bring together environmental factors with equine strongyle diversity in Ukraine concluded that foals and horses older than 16 years displayed higher species richness (Kuzmina et al., 2016). However, species richness is only a limited facet of the actual biodiversity as it does not account for the relative proportions of every species found. Further, the impact of host sex was not considered (Kuzmina et al., 2016).

Host sex effect is a major driver of helminth community assemblage, male mammals being generally more frequently and more heavily parasitized (Poulin, 1996; Zuk and McKean, 1996). Although, some discrepancies exist to this general pattern (Grzybek et al., 2015). In horses, no clear consensus has been reached so far. The analysis of necropsy reports in Australia found differential prevalence and abundance of 13 endoparasites between the two sexes (Bucknell et al., 1995). However, no general trend emerged as females were more heavily infected by *Cyathostomum pateratum* but less frequently infected by *Cylicodontophorus bicoronatus, Cylicocyclus insigne* and *Cylicocyclus elongatus* than stallions and geldings (Bucknell et al., 1995). Study of faecal egg count in a feral horse population also reported a blurred pattern between the two sexes as the polarity of the sex effect was spatially contrasted (Debeffe et al., 2016). Other studies reported higher faecal egg count in geldings or could not evidence any difference (Kornaś et al., 2010; Kornaś et al., 2015).

Analysing strongyle communities recovered from 48 horses in Poland, this study proposes to investigate species community structure and diversity, and assess how host sex drives these.

## 2. Material and methods

### 2.1. Study population

Strongyles were collected from 48 horses scattered across five riding schools and one stud farm from Poland. The study population was composed of 37 mares and 11 stallions. The Małopolska breed accounted for most of the sampled horses (n=25), in addition to some Hucul (n=6) and pure blood Arabian horses (n=17) that were found in only two sites (supplementary figure 1). Two third of the individuals were born before 2007 (supplementary figure 1), horses sampled in riding-schools being 6 years older than those from stud farms on average.

Within each premise, the number of sampled horses ranged from 3 to 18 individuals (Table 1, supplementary figure 1).

**Table 1.** Premise features and ecological properties of recovered strongyle communities. This table shows the number of sampled horses per sex and site (SF: Stud farm; RS: Riding school). Average species richness, *i.e*. number of recovered strongyle species and Shannon’s index of diversity are also provided by premise. Bottom row provided overall total number of horses and average species richness and Shannon’s index.

| Site | Type | No. horses | Mare | Stallion | Species richness | Shannon index |
| --- | --- | --- | --- | --- | --- | --- |
| 1 | Stud farm | 18 | 18 | 0 | 12.17 (2.04) | 2.07 (0.17) |
| 2 | Riding school | 14 | 8 | 4 | 9.86 (3.74) | 1.59 (0.48) |
| 3 | Riding school | 3 | 2 | 1 | 6.00 (0) | 1.46 (0.23) |
| 4 | Riding school | 3 | 1 | 2 | 6.67 (1.53) | 1.43 (0.12) |
| 5 | Riding school | 3 | 0 | 3 | 11.67 (6.51) | 1.75 (0.38) |
| 6 | Riding school | 7 | 6 | 1 | 11.00 (2.16) | 1.90 (0.14) |
|  |  | 48 | 35 | 11 | 10.6 (3.4) | 1.8 (0.38) |

Parasite control strategy in every location relied on two ivermectin treatments performed twice a year in March and October, before and after pasture season respectively.

### 2.2. Strongyle collection and identification procedure

The species composition of strongylid nematodes was performed by collecting 500g of faecal material expelled 24 hours after ivermectin treatment following the strategy of other trials (Osterman Lind et al., 2003; Kuzmina et al., 2005). Subsequently, samples were rinsed with tap water on 250μm mesh sieves and examined under stereomicroscope for the presence of strongylids. Collected nematodes were fixed in 70% ethyl alcohol with 5% glycerine additive to clarify their body structure. Strongylids were identified according to previously published morphological keys (Lichtenfels et al., 2008). For the sake of clarity, strongyle species names have been abbreviated to a three-letter identifier throughout the manuscript. Table of correspondence has been provided as supplementary information (Table S1).

Faecal egg count before treatment was performed on 5g of faecal material, diluted in 70 mL of a flotation solution (NaCl, d=1.18), with a sensitivity of 50 eggs/g.

### 2.3. Community structure analyses

Statistical analyses were implemented with R v 3.4.1 (R Core Team, 2016), and diversity analyses were run using the vegan package v. 2.4-3 (Oksanen et al., 2017). Strongyles recovered from every horse were hereafter referred to as a community. Within each community, species intensity referred to the number of worms recovered from a given horse, while species relative abundance in a given community corresponded to the proportion of worms found relative to the total worm burden. Species prevalence was defined at the premise level as the proportion of horses infected by a given strongyle species.

Species richness was defined as the total number of species recovered from each horse. Species diversity within each horse, corresponding to Whittaker’s α-diversity (Witthaker, 1960), was measured by the Shannon’s index (Shannon, 1949) defined as:

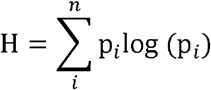

where *p_i_* is the proportional abundance of species *i*.

Species turnover can occur along environmental gradients, hence resulting in variation among individual communities from a given group. This measure of dissimilarities between communities was originally coined as β-diversity (Witthaker, 1960), and will contribute only marginally to the global diversity if individual communities show little or no variation among groups. Difference in β-diversity among environmental factors (location and horse sex) was tested following a multivariate framework (Anderson, 2006; Anderson et al., 2006). This framework measures β-diversity as the distance of every community to the average group community, determined as the centroid of a principal component analysis of community structures (Anderson et al., 2006).

Non-random species co-occurrence was tested following a probabilistic framework that compares observed species frequencies to the expected frequencies under species independence (discarding species pairs when the expected number of co-occurrence was less than 1) (Veech, 2013; Griffith et al., 2016). Species interactions could also result in positive or negative feedbacks that could impact on species relative abundances. To test for these interactions, species counts were regressed upon other species intensity in a pair-wise manner after accounting for environmental variables (horse sex and sampling site) and by the mean of a Poisson regression. For this analysis, the 13 species with more than 200 individuals across horses were considered, as failure to do so results in strong positive associations for the minor present species. Because of the high number of statistical tests (n=105), a Bonferroni correction was applied and a p-value below 0.05/105=5x10^-4^ was considered significant.

A Shapiro-Wilk test was run to test the normality of species richness. Pearson’s correlations were computed using the rcorr() function of the Hmisc package v.4.0-3 (Harrell and Dupont, 2017) and visualized as a network with the GGally package v.1.3.2 (Schloerke et al., 2017).

### 2.4. Host sex effect on species relative abundance and prevalence

The impact of host sex on strongyle abundance (worm counts) and prevalence was investigated for the 11 core strongyle species, *i.e*. most abundant and most prevalent. *C. catinatum* (CAT) was the best disperser (prevalence of 97.9%) and was not included in the prevalence data analysis as not enough variability was present to assess factors of variation. To account for the data structure (supplementary figure 1), these effects were investigated using the only data from premises where horses of each gender were present (premises 2, 3, 4 and 6; n=27 horses).

Worm counts and species prevalence were modelled using a negative binomial and logistic regression respectively, fitting the sampling site, strongyle species, horse sex and their interaction as fixed effects as follows:

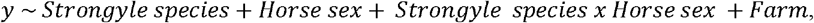

where y stands for species count or prevalence accordingly.

A linear regression analysis was applied to Shannon’s index values to evaluate variation between farms and sexes.

For every model, the sex effect on species relative abundance and prevalence was estimated across farms (19 mares and 8 stallions). The farm effect accounted for both management and breed type differences.

## 3. Results

### 3.1. Strongyle dispersal abilities were contrasted and inversely correlated with their fecundity

On average, 608.7 worms per horse were collected, ranging from 111 to 2,884 (supplementary table 2). Overall, 23 species were found across premises (supplementary table 2), with average species richness ranging from 6 to 12 ± 2 (Table 1). Between four and 20 species could be recovered at the horse level (supplementary table 2). Total worm count was not significantly correlated with pre-treatment faecal egg count (Spearman’s *r* = 0.22, p=0.13).

CAT and BIC were respectively the most and the least abundant species, worms of each species summing up to 5,462 and one worms recovered across communities. *S. vulgaris* was encountered in only 6% of horses. Relative species abundance across communities could distinguish between two groups of species, being either highly abundant and prevalent (Figure 1A) or with lower dispersal ability and minor contribution to communities (figure 1A). CAT, NAS and LON were found in more than 75% of horses and accounted for at least 10% of recovered worms (Figure 1A).

**Figure 1.**
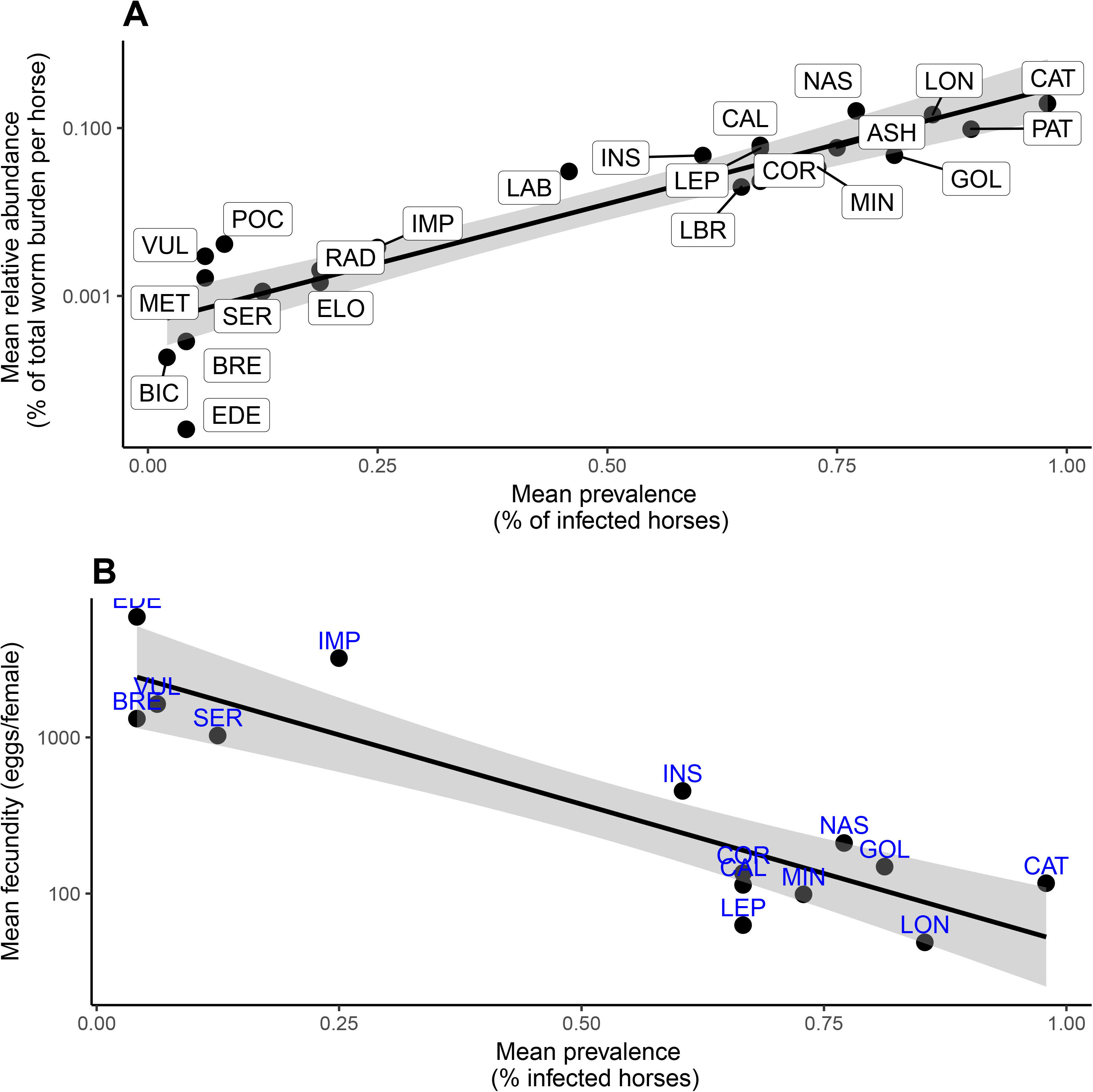
Dispersal ability of strongyle species. Figure 1A represents the relationship between species relative abundance and its prevalence across sites and seasons. Figure 1B displays the relationship between species prevalence and its published fecundity in eggs per female. In each case, regression line and 95% confidence interval (grey area) is represented.

Pearson’s correlations between worm fecundity and species abundance and prevalence were estimated to determine whether fecundity could underpin better dispersal abilities. These correlations suggested that fecundity was negatively correlated with species prevalence (*r*=-0.71, p=0.004; figure 1B) and that the same trend held for species relative abundance (*r*=-0.48, p=0.08; supplementary figure 2).

Species richness accumulation curves were generated for the two farms with biggest herd size to estimate how much sampling effort should be made to sample community diversity. These curves suggested that the minimal sampling effort had to be comprised between seven and nine horses to detect 90% of total species present in one location (figure 2).

**Figure 2.**
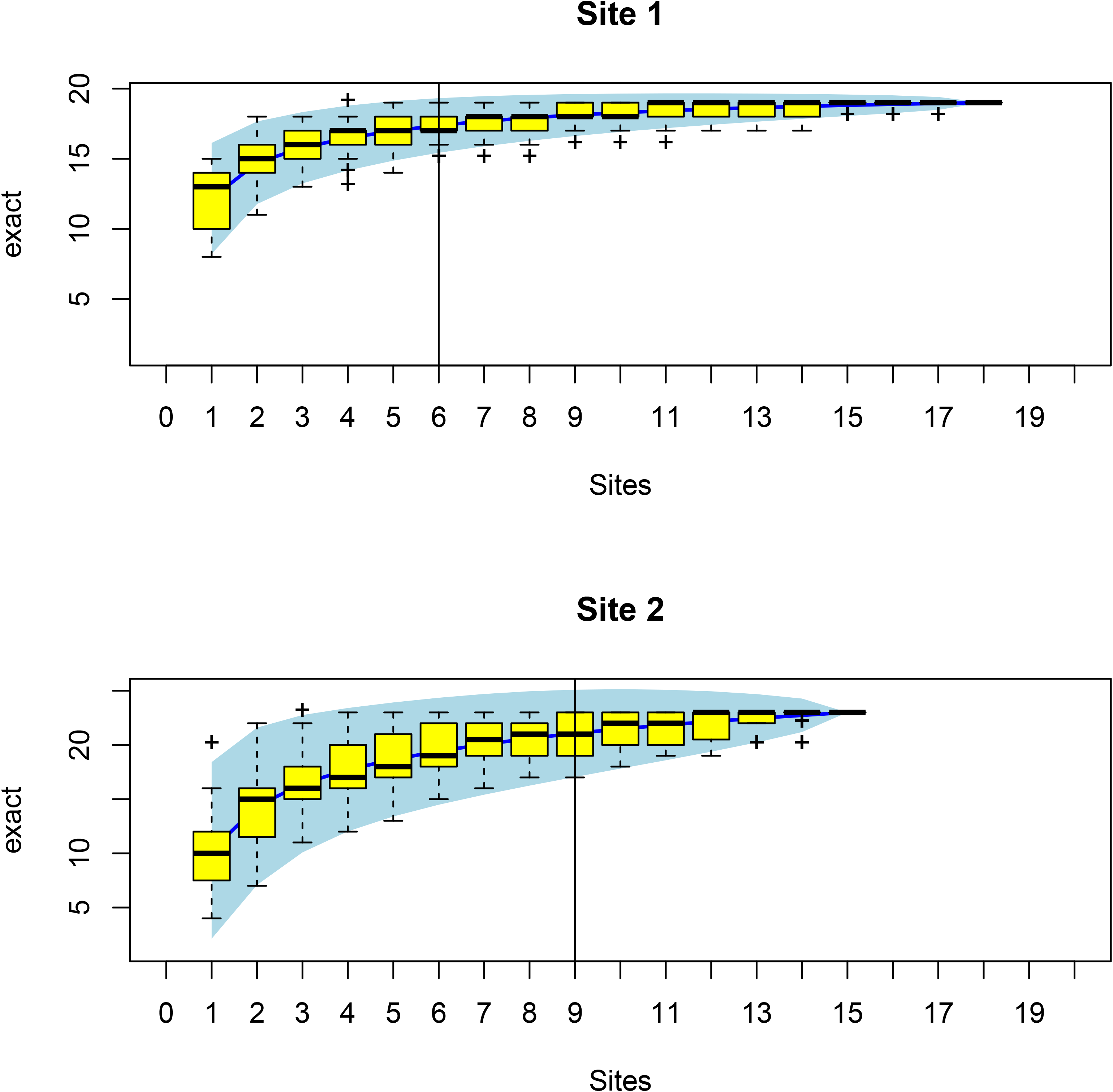
Species accumulation curves for sites one and two. For these two sites, the sampling effort (x-axis) required to detect a given number of species (y-axis) is plotted as a blue line with 95% confidence interval (blue area). Box-plots stand for random variation measured by permutations. Vertical lines assess the sampling effort able to detect to 90% of species in a given site.

### 3.2. Communities were structured by a network of mostly positive interactions

Species co-occurrence analysis was dominated by random co-occurrences that represented 87% of the 200 tested combinations (Figure 3A, supplementary table 3). Nonetheless, a few species pairs seemed to positively interact with each other (n=26, figure 3A, supplementary table 3), including some core species, *e.g*. LON-NAS (p=0.04) or PAT-GOL (p=0.04).

**Figure 3.**
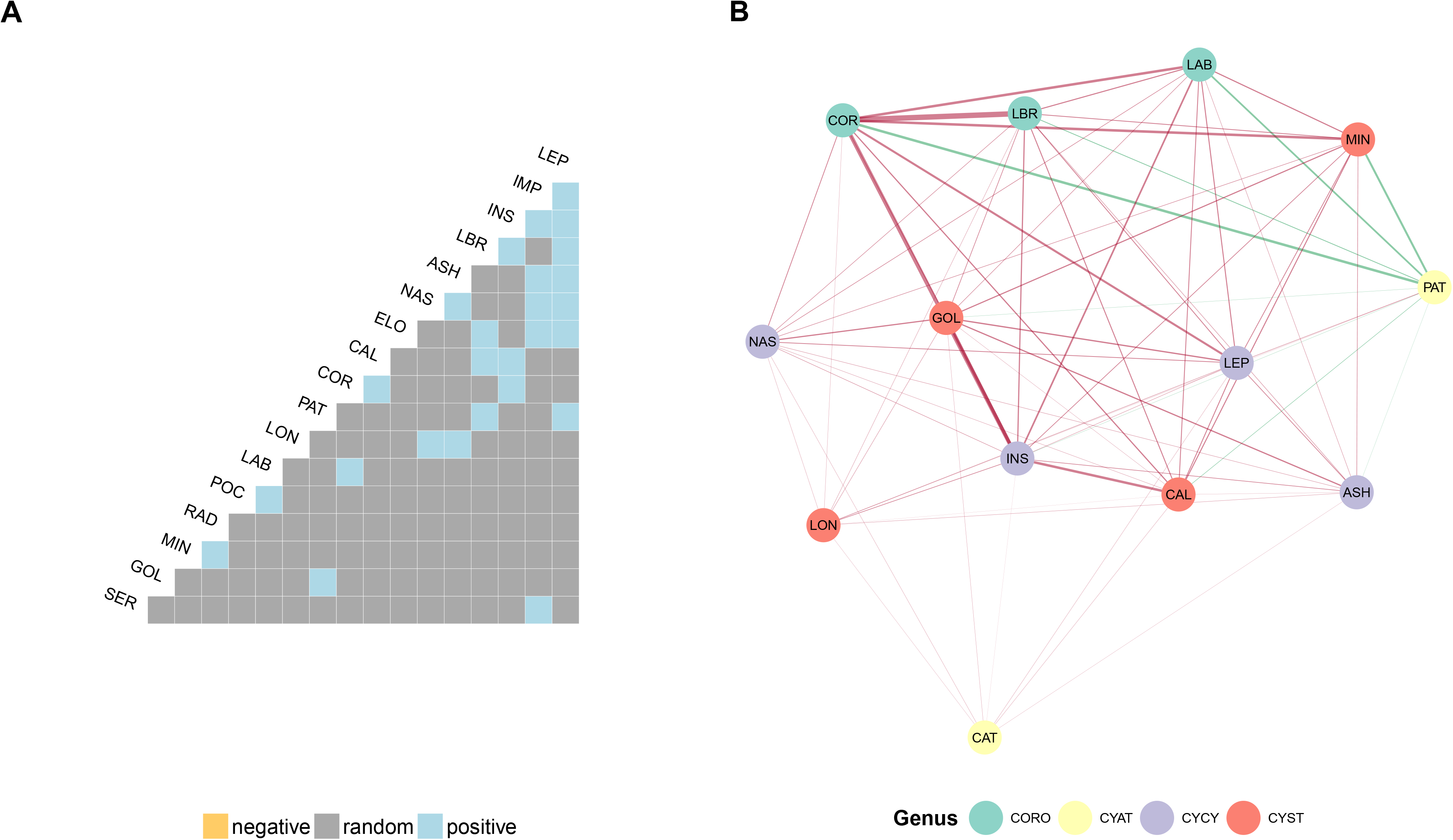
Species co-occurrence and interactions network based on relative abundances for recovered strongylid species. Figure 3A displays a pair-wise matrix of co-occurrence between species co-occurring more than once. Grey boxes indicate that species pairs occurred randomly, whereas 26 pairs co-occurred significantly more often than expected. Figure 3B represents a network, drawn using a force-directed algorithm from the matrix of interactions estimated by pair-wise Poisson regression on species counts after accounting for environmental effects. To avoid spurious positive associations due to low counts, the analysis was restricted to the only species totalizing 200 worms overall. Nodes act as repelling objects that are organized to minimize forces in the network. Each node represents a species and is coloured according to the genus it belongs to (CORO: *Coronocyclus;* CYAT: *Cyathostomum;* CYST: *Cylicostephanus;* CYCY: *Cylicocyclus*). Connected nodes entertain significant association after Bonferroni correction for multiple testing (p<6x10^−4^). Edges are coloured to reflect the correlation sign (red if positive, green if negative) and their widths mirror the association intensity (the thicker, the bigger absolute effect). For clarity purpose, species names have been abbreviated and can be found in supplementary table 1.

On the top of the presence/absence covariation analysis, species intensities were regressed upon each other after correcting for environmental factors, to evaluate how the presence of one species could positively or negatively impact on the others. Significant interactions were used to build a network between *Coronocyclus* (CORO), *Cyathostomum* (CYAT), *Cylicostephanus* (CYST) and *Cylicocyclus* (CYCY) species (Figure 3B) that was also mostly driven by positive interactions between species intensities (Figure 3B). Notably, PAT was the only species whose intensity varied negatively with other species, with the exception of LON and LEP (Figure 3B). Three of the best dispersers (CAT, NAS, LON) seated on the edge of the network, with weak links to other species, suggesting that variation in their intensity was not impacted by other species. On the contrary, *Coronocyclus* species, *i.e*. COR, LBR and LAB, were more tightly connected, suggesting similar behaviors.

### 3.3. Horse gender affects dissimilarity between strongyles communities and species abundance

Strongyle community diversity ranged from 0.6 to 2.6 at the horse level, but was more consistent at the farm level with average values comprised between 1.43 ± 0.12 and 2.07 ± 0.17 (Table 1). No significant variation in species alpha-diversity was found between farms (p=0.2).

Neither did horse sex significantly affect community diversity (p=0.88), suggesting alpha-diversity was equivalent between mares and stallions.

An analysis of community dissimilarity was performed to assess the extent of species turnover across horse gender and sampling sites. Strongyle communities structure significantly varied between sampling sites (p=0.02) and were also significantly less homogeneous in mares than in stallions (p=0.003).

No systematic horse sex driven differences in strongyle species abundance was identified across strongyle species by the binomial regression approach (p=0.57, Table 2). However, a significant interaction was found between horse gender and strongyle species abundance. Notably, CAL was significantly more abundant in stallions than in mares, whereas the opposite trend (non-significant) was generally found for other species (figure 4, supplementary table 4). This pattern was also conserved when comparing CAL abundance between site 1 (14.8 worms/horse on average; mares only) and site 5 (107 worms/horse on average; stallions only) even though this difference could not be disentangled from a site-associated effect.

**Figure 4.**
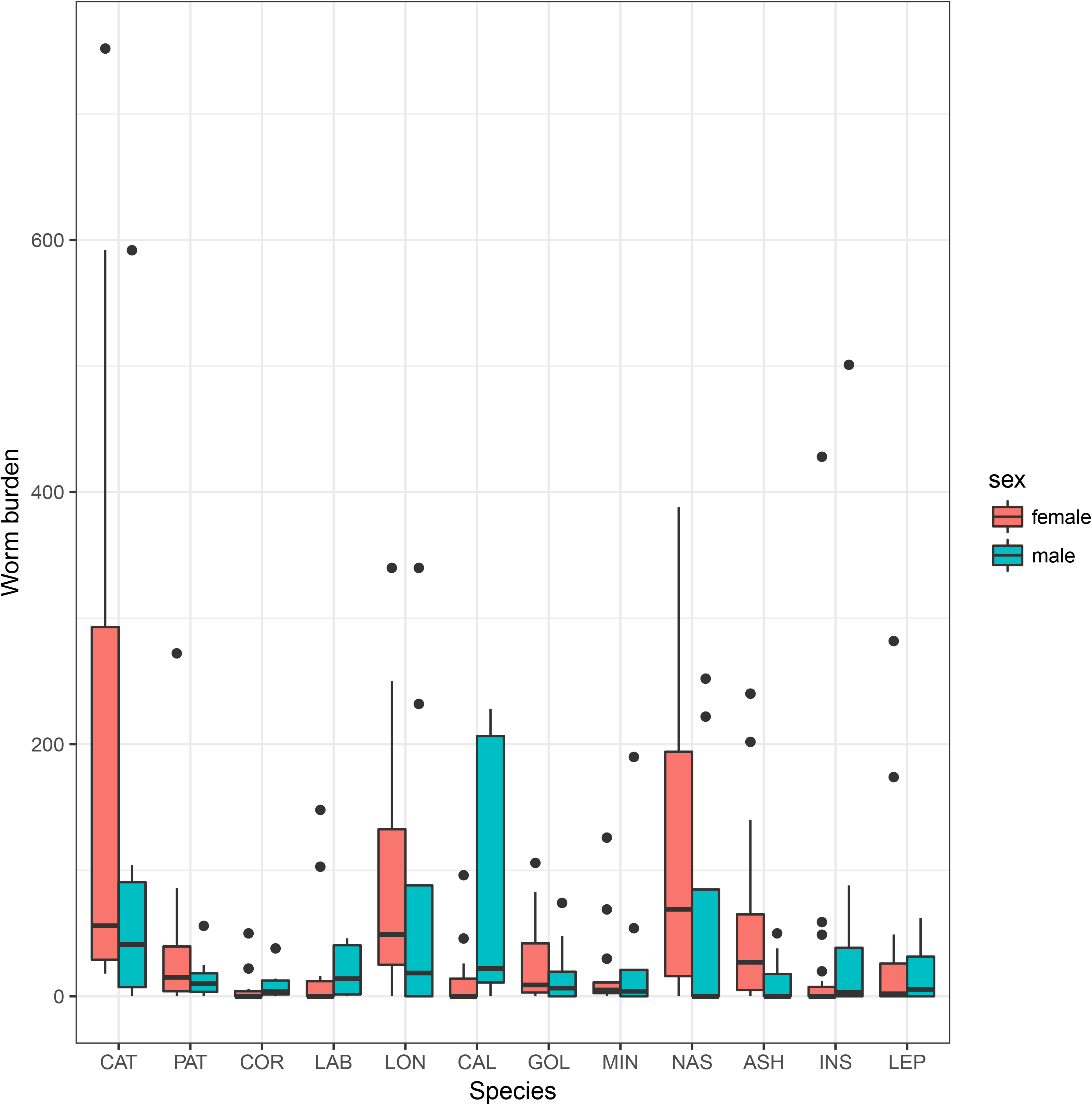
Strongyle abundance variation according to horse gender. Boxplots of strongyle counts are plotted against horse gender (red for females and blue for males) across farms. Of note, is the higher abundance for *Cylicostephanus calicatus* (CAL) in males. For clarity purpose, species names have been abbreviated and can be found in supplementary table 1.

Strongyle dispersal measured by the prevalence of each species did not significantly vary between mares and stallions, although NAS was found slightly more often in stallions (p=0.06, supplementary table 4).

## 4. Discussion

This study focused on the structure of equine strongyle communities under eastern Europe climatic conditions. Data suggested functional trade-offs between species dispersal ability, measured by prevalence and relative abundance, and species fecundity that may explain species coexistence. In addition, horse sex seems to be a significant driver of strongyle community structure and diversity.

Numerous reports on equine strongyle community structure have already evidenced the bimodal pattern of their assemblage, with five to ten highly prevalent and abundant species and additional minor species (Ogbourne, 1976; Mfitilodze and Hutchinson, 1990; Lyons et al., 1999; Collobert-Laugier et al., 2002; Kuzmina et al., 2005; Traversa et al., 2010). Our data also highlighted the same split between core and satellite species (Hanski, 1982), with a set of three successful dispersers, *i.e. C. catinatum, C. nassatus* and *C. longibursatus*, that were found in at least 75% of horses and represented at least 10% of the total worm burden. Not only corroborating the known literature, this finding also provided good support for the chosen sampling strategy (Osterman Lind et al., 2003; Kuzmina et al., 2005). A single sampling 24 hours after treatment might have biased the sampling efficiency downwards as it has been suggested that a significant proportion of parasitic material could still be recovered after 36 and up to 60 hours (Kuzmina et al., 2005). From our results, some species reported as minor in the present study (*S. edentatus*, *S. vulgaris*, *T. serratus* and *C. elongatus*) may have suffered from the sampling strategy as most of them were expelled 48h after treatment onwards in a previous trial (Kuzmina et al., 2005). Notably, this prevents any firm conclusion regarding the minor contribution of *S. vulgaris* to the strongyle communities, even though displayed prevalence level were in agreement with other reports from regularly dewormed horses (Lind et al., 1999; Boxell et al., 2004). Nevertheless, community structure and species richness estimates were similar to previous reports under eastern European conditions (Kuzmina et al., 2005; Kuzmina and Kharchenko, 2008), ruling out an overall down sampling of the actual diversity. In addition, *C. coronatus* was among the most frequently encountered species whereas it usually occurs in the upper portion of the large intestine (Collobert-Laugier et al., 2002; Stancampiano et al., 2010), supporting the unbiased community structure reported herein.

Under the considered conditions, a limited number of horses were required to sample 90% of the strongyle community diversity. This is of particular relevance for the implementation of field sampling where necropsy is not an option and minimal interference with premise activity are expected.

Although the overall structure of strongyle community has been well characterized in equids (Ogbourne, 1976; Mfitilodze and Hutchinson, 1990; Lyons et al., 1999; Kuzmina et al., 2005; Traversa et al., 2010), little is known about the mechanisms and variation factors underpinning its stability. Remarkably, sampled communities were entertaining positive interactions, both in terms of presence/absence patterns or in their intensity covariation. This finding was in contradiction with previously reported results from an abattoir survey, that highlighted negative interactions between major species (Stancampiano et al., 2010). Although, the negative interactions mostly occurred between species that were less represented in our present report, *i.e. S. vulgaris* and *S. edentatus* or *Triodontophorus* species (Stancampiano et al., 2010). But results from the previous survey were grounded on relatively low infection intensity as only 1,590 adult worms were found from 60 horses (26.5 worms per horse on average) (Stancampiano et al., 2010). On the contrary, horses sampled in the current study yielded an average of 538 adult worms per horse, providing enough resolution to interrogate species interactions. In addition, our analyses were restricted to the best dispersers, hence preventing spurious positive associations that can occur when correlating low counts (Poulin, 2001). Our results were in line with known positive associations between core species (Holmes, 1987). Reports of helminths species co-occurring more often than expected by chance have also been made for human and rodent systems (Behnke, 2008). These positive interactions could arise from niche partitioning between species within horses. It could explain positive feedbacks between *Cylicostephanus goldi* and *C. nassatus* for instance that mostly occupy the dorsal and ventral colon respectively (Collobert-Laugier et al., 2002; Stancampiano et al., 2010), but does not support positive interactions between *Cylicostephanus minutus* and *Cylicostephanus calicatus* that tend to share the same organs (Collobert-Laugier et al., 2002; Stancampiano et al., 2010). It may hence suggest that some of the equine strongyle species share mutualistic interactions as already reported for rodent models (Behnke, 2008).

But the existence of functional trade-offs could also explain this pattern. Indeed, observations from plants (Angert et al., 2009) or insects (Duthie et al., 2015) highlighted that negative environmental impact on species competitive ability could favour their coexistence. Empirical data from fig wasps under a fluctuating environment demonstrated a dispersal-fecundity trade-off, the more fecund individuals occurring in resource-dense area (Duthie et al., 2015). In such a fluctuating environment where resource availability can vary, competition may be erased as the best competitors are not able to colonize every available niche. The negative correlation between known equine strongyle fecundity (Kuzmina et al., 2012) with their observed dispersal capacities reported herein may also suggest that the same kind of trade-off exists, ultimately leading to species coexistence. Last, the negative interactions that *C. pateratum* entertained with other species remains unexplained and could sign exploitation (space, nutrient) or interference (local inflammation, cross-reactivity) competition (Bottomley et al., 2007).

Host sex is a significant driver of parasite community, males being generally more heavily infected than females (Poulin, 1996). This pattern does not always hold (Grzybek et al., 2015), especially in horses where definitive consensus has not been reached yet (Kornaś et al., 2010; Kornaś et al., 2015; Debeffe et al., 2016). Our results suggested that horse sex did not exerted a conserved influence on worm relative abundance across species, but that mares had more consistent strongyle communities. In addition, higher *C. calicatus* worm counts were found in stallions. This finding was already reported in a previous Australian survey, that also evidenced additional host sex interactions for *C. coronatus* and *C. catinatum* (Bucknell et al., 1995). Variation in the qualitative composition of strongyle structure might hence contribute to the lack of clear difference between male and female excretion patterns. But mechanistic explanations to support a higher turnover in males remains unclear and deserve further exploration. In the case of species-bias in anthelmintic resistance, this could favour the implementation of differential management between the two sexes.

## Conclusions

This survey provided insights into applied and more fundamental knowledge about equine strongyle communities and how host sex drives their structure and diversity. The bimodal structure of equine strongyle communities split into core and satellite species was confirmed, underscoring the overwhelming contribution of *C. nassatus, C. catinatum* and *C. longibursatus*. We could also evidence that less than ten horses were needed to sample most of the diversity, which is particularly relevant for logistically constrained surveys.

More fundamentally, it seems that equine strongyle communities are largely structured around a network of positive, if at all, interactions, with the only exception of *C. pateratum*. While mutualistic interactions and niche partitioning could explain this pattern, the observed trade-offs between fecundity and dispersal abilities may also explain this coexistence.

Horse sex did not appear to influence the overall worm counts, but a higher species turnover was found among stallions. This latter finding would be particularly relevant in case of differential resistance to anthelmintics between species, that would eventually lead to contrasted recommendations between mares and stallions.

While our data provided new insights into the structuring equine strongyle communities, only a limited set of the potential factors shaping these communities have been tackled. Further research in this area should make use of recent advances in sequencing technology to increase the sensitivity of worm sampling and also interrogate putative interactions with the host gut microbiota.

## Acknowledgements

The authors are particularly indebted to the premise managers who took part to this survey.

---

**Supplementary Figure 1.**
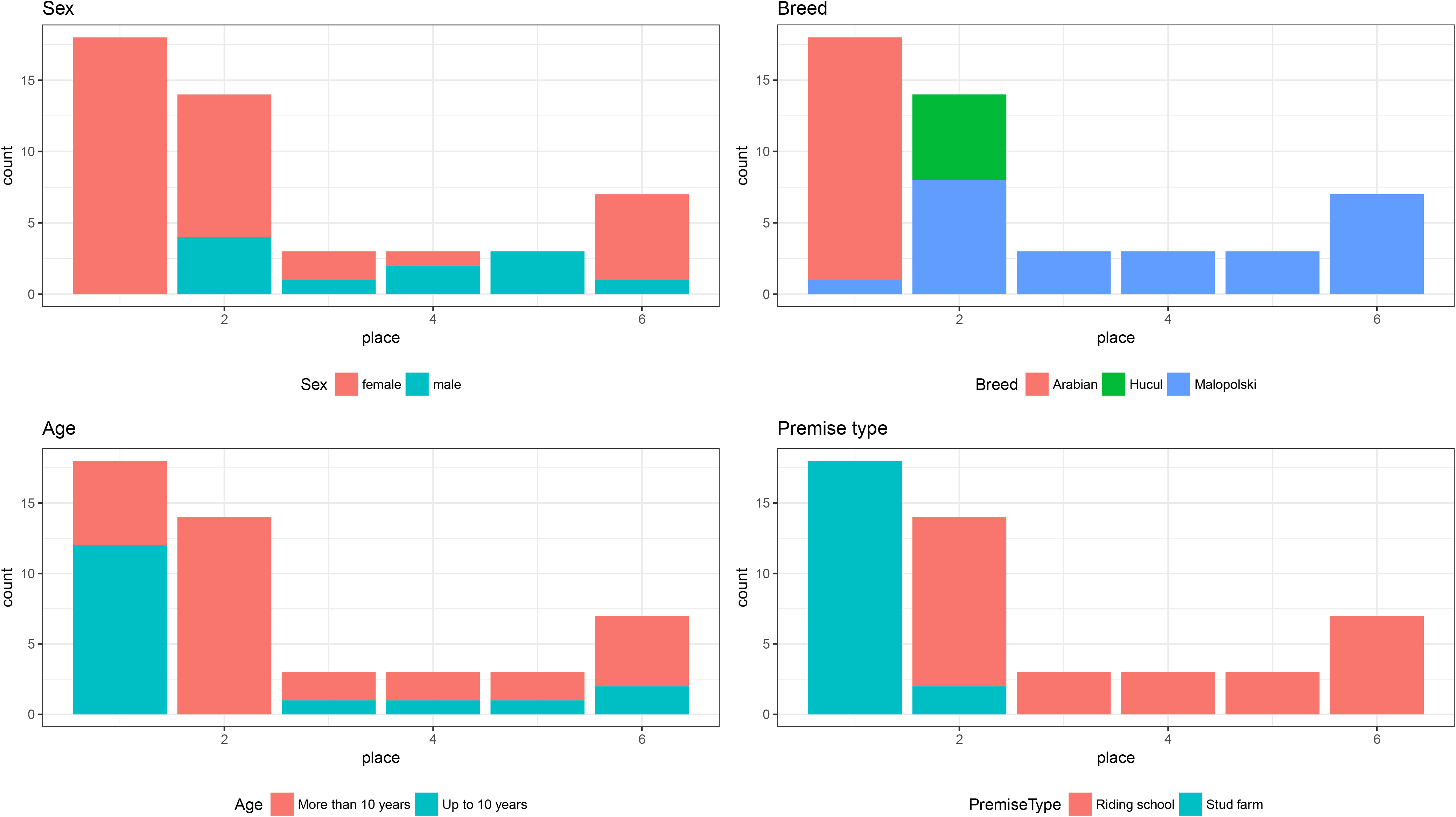
Full data structure

**Supplementary Figure 2.**
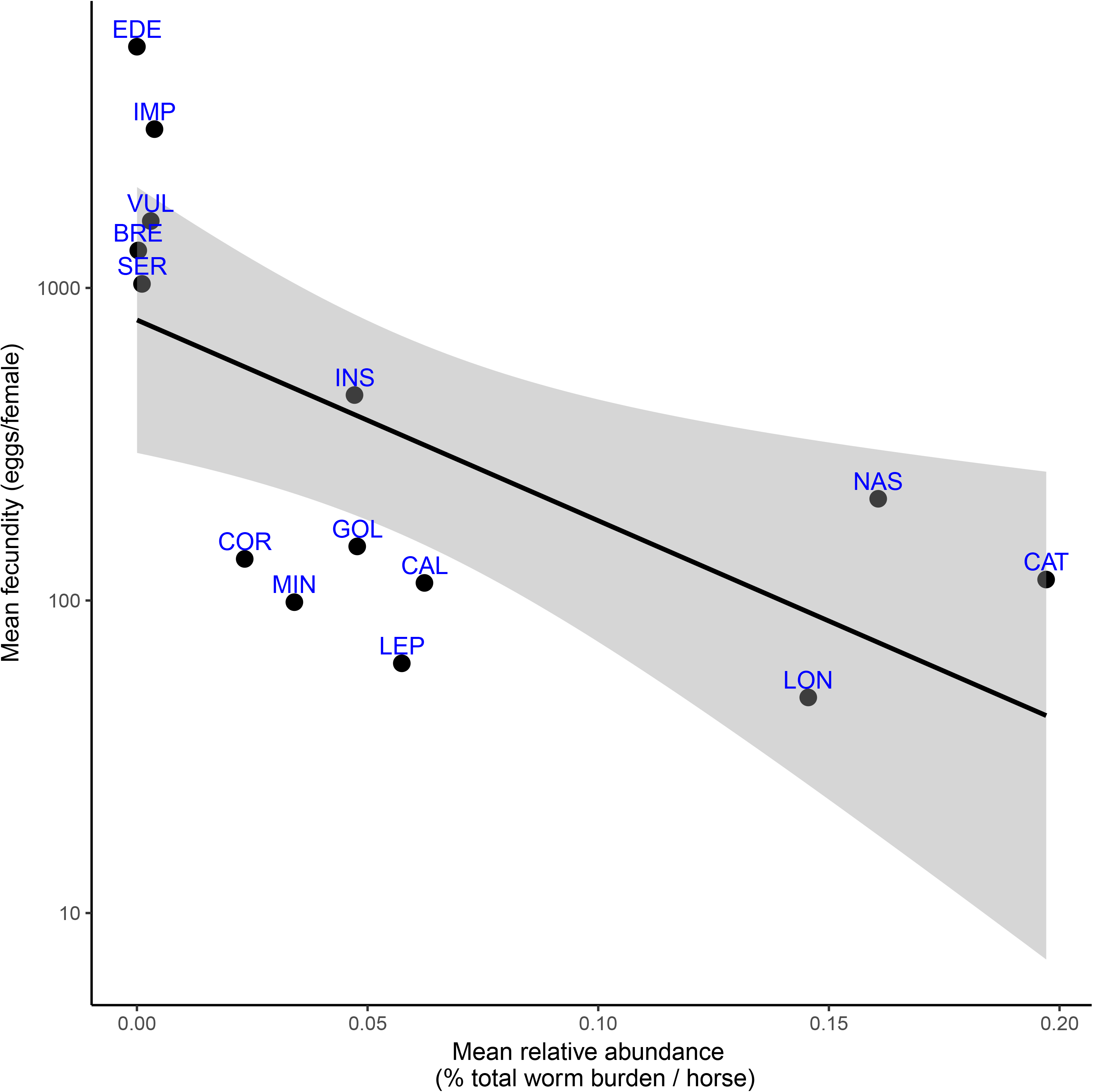
Relationship between species relative abundance and fecundity

**Supplementary Table 1.** Recovered species names and corresponding abbreviation

**Supplementary Table 2.** Individual worm counts and associated metadata

**Supplementary Table 3.** Table of co-occurrence probabilities between species

**Supplementary Table 4.** Pair-wise regression coefficients between the 13 most abundant species

**Supplementary Table 5.** Species intensity and prevalence modelling output

